# Interaction of adenylate cyclase CyaC with regulators Smc01817 and Clr of *Sinorhizobium meliloti*: cAMP as an ON/OFF trigger in sensor-regulator complexes?

**DOI:** 10.1101/2024.01.22.576614

**Authors:** Robin Klein, Juliane Wissig, Katharina Küllmer, Gottfried Unden

## Abstract

The symbiotic bacterium *Sinorhizobium meliloti* contains a large number of adenylate cyclases (AC) for the control of different life styles. ACs produce cyclic AMP (cAMP) as a secondary messenger. Earlier, the redox responsive membrane-intrinsic AC CyaC has been shown to produce a trimeric signalling complex CyaC×CycR×cAMP-Clr. Here, use of the bacterial two-hybrid system BACTH showed that the LysR-type transcriptional regulator Smc01817 interacts *in vivo* with CyaC and the general cAMP-regulator Clr, suggesting the formation of a sensor-regulator complex CyaC×Smc01817×cAMP-Clr. Therefore, formation of sensor-regulator complexes from ACs and transcriptional regulators seems to be a means in *S. meliloti* to setup specific signalling routes in a background with a large number of signalling routes applying the same signalling molecule (cAMP). Use of caged cAMP allows to address a specific signalling route and set of regulators, which is at variance with the classical role of cAMP as a diffusing secondary messenger.

## Introduction

*Sinorhizobium meliloti* is characterized by a complex life style including free-living and symbiotic growth. Adaptation to the different forms of growth requires major changes in physiology, cell composition and the capacity to interact with plant components and depends on multiple sensing and regulatory processes: The bacteria are able to respond to a large number of growth parameters, including the availability of electron acceptors, the carbon and nitrogen source, and physical and chemical stimuli provided from the environment and the plant host (Masson et al. 2009; Tian et al. 2012; Zou et al. 2017). A large set of genes is required for adaptation and the developmental process, which are controlled by a wide range of sensory and signalling devices. Among others, a large number of adenylate cyclases are employed in the adaptation process from free living to symbiotic life-style. Up to 28 adenylate cyclases have been identified in *S. meliloti* (Capela et al. 2001; Krol et al. 2016) which are supposed to sense environmental cues and adapt the bacterial cell to the metabolic changes.

Co-existence and specific function of a large number of sensory/signalling devices relying on the same signalling molecule cyclic AMP (Capela et al. 2001; Krol et al. 2016) requires specific organization and set-up of the signalling pathways to ensure that a potentially freely diffusing molecule such as cAMP activates one pathway but not another. Recently, the signalling pathway starting from adenylate cyclase CyaC has been shown to form ternary sensor-regulator complexes that can present the prerequisite for specific signalling from the sensory to the regulatory site. CyaC represents a membrane-intrinsic class III AC (Krol et al. 2016; Linder & Schultz 2003). Thus, CyaC forms a ternary complex with a specific (CycR) and an overriding transcriptional regulator (Clr). The trimeric sensor-regulator complex CyaC×cAMP-Clr×CycR is formed upon activation of the adenylate cyclase, connected to the reaction of ATP and the formation of cAMP (Klein et al. 2023). Formation of the complex is apparently linked to the binding of cAMP. The ‘caged’ cAMP forms a CycR×cAMP-Clr sub-complex which is supposed to bind after release from the adenylate cyclase to target promoters. In the dimeric complex the specific regulator CycR is supposed to bind together with the general cAMP binding transcriptional regulator Clr to promoters to convey specific and cAMP mediated control. The complex formation between CyaC, CycR and Clr has been shown in *in vivo* studies using the bacterial two-hybrid system in *E. coli* (Karimova et al. 1998, 2005; Klein et al. 2023) and was verified *in vitro* with purified proteins. The specific transcriptional regulator CycR of the system is encoded by the *cycR* (or *smc01819*) gene located directly downstream of the gene encoding the adenylate cyclase (*cyaC*, or *smc01818*) (Fig. 1). Remarkably, the *smc01817* gene located upstream of *cyaC* encodes a transcriptional regulator of the LysR type (Luo et al. 2005). LysR-type transcriptional often consist of N-terminal DNA-binding and C-terminal effector-binding domain (Schell 1993; Maddocks and Oyston 2008) which are conserved in the Smc01817 protein. Given the frequent clustering of functionally related genes, it was tested here by *in vivo* BACTH studies, whether there are indications for the interaction of Smp01817 with CyaC and Clr, similar to the trimeric CyaC×cAMP-Clr×CycR complex.

## Materials and Methods

### Bacterial strains, plasmids BACTH assay

The plasmids used are listed in Table 1. The molecular genetic methods, including cloning, generation of gene fusions, DNA isolation, and manipulations were performed according to standard procedures (Sambrook et al. 1989; Miller 1992) as described in Klein et al. (2023). Bacteria were grown aerobically or anaerobically at 37°C or 30°C (Kim et al. Klein et al. 2023). The BACTH assays (Karimova et al. 1998) were performed in strain *E. coli* BTH101(F-*cya-99, araD139, galE15, galK16, rpsL1, hsdR2, mcrA1, mcrB1* (StrR)) transformed with the respective plasmids encoding the fusion proteins with T18 and T25 fragments of 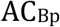. The proteins were cloned in plasmids pUT18, pUT18C, pKT25 and pKNT25 for producing the T18 and T25 fragment for C- and N-terminal fusion, respectively (Klein et al. 2023). For the BATCH assays, the bacteria were grown anaerobically in LB broth with 20 mM DMSO which produces highest activities (Wissig et al. 2019) in microtiter plates to an OD578 0.6 to 0.9. The β-galactosidase assays (Monzel et al. 2023; Wörner et al. 2016) were performed as described (Klein et al. 2023) and are presented as the mean (with standard deviation) from at least two biological and four technical replicates each.

**Table 1.** Plasmids used.

| **Plasmid** | **Genotype** | **Reference** |
| --- | --- | --- |
| **pKT25** | pSU40, for C-terminal T25 fusion, Km<sup>R</sup> | Karimova et al. 2005 |
| **pKNT25** | pSU40, for N-terminal T25 fusion, Km<sup>R</sup> | Karimova et al. 2005 |
| **pUT18** | pUC19, for N-terminal T18 fusion, Ap<sup>R</sup> | Karimova et al. 2005 |
| **pUT18C** | pUC19, for C-terminal T18 fusion, Ap<sup>R</sup> | Karimova et al. 2005 |
| **pKT25-zip** | pKT25, but T25-zip, Km<sup>R</sup> | Karimova et al. 2005 |
| **pUT18C-zip** | pUT18C, but T18-zip, Ap<sup>R</sup> | Karimova et al. 2005 |
| **pMW689** | His<sub>6</sub>-pUT18, but CyaC(K410/T484A)-T18, Ap<sup>R</sup> | Klein et al. 2023 |
| **pMW692** | His<sub>6</sub>-pKNT25, CyaC(K410/T484A)-T25, Km<sup>R</sup> | Klein et al. 2023 |
| **pMW3087** | pUT18, but Clr-T18, Ap<sup>R</sup> | Klein et al. 2023 |
| **pMW3088** | pUT18C, but T18-Clr, Ap<sup>R</sup> | Klein et al. 2023 |
| **pMW3089** | pKT25, but T25-Clr, Km<sup>R</sup> | This work |
| **pMW3090** | pKNT25, but Clr-T25, Km<sup>R</sup> | Klein et al. 2023 |
| **pMW2868** | His<sub>6</sub>-pKT25, but T25-Smc01817, Km<sup>R</sup> | This work |
| **pMW2869** | His<sub>6</sub>-pKNT25, but Smc01817-T25, Km<sup>R</sup> | This work |
| **pMW2870** | His<sub>6</sub>-pUT18, but Smc01817-T18, Ap<sup>R</sup> | This work |
| **pMW2871** | His<sub>6</sub>-pUT18C, but T18-Smc01817, Ap<sup>R</sup> | This work |
| **pMW1424** | pUT18C, but T18-NreC (*S. carnosus*), Ap<sup>R</sup> | Niemann et al. 2014 |
| **pMW1439** | pUT18, but NreC-T18 (*S. carnosus*), Ap<sup>R</sup> | Niemann et al. 2014 |
| **pMW1777** | pKNT25, but NreC-T25 (*S. carnosus*), Km<sup>R</sup> | Niemann et al. 2014 |
| **pMW1780** | pKT25, but T25-NreC (*S. carnosus*), Km<sup>R</sup> | Niemann et al. 2014 |

## Results

### Interaction of CyaC with Smc01817

To evaluate the presence of an adenylate cyclase – regulator complex of the CyaC×CycR×Clr type, interaction of the suspicious regulator Smc01817 with adenylate cyclase CyaC and the general transcriptional activator Clr was tested *in vivo* by the use of the bacterial adenylate cyclase based two-hybrid system (BACTH). CyaC and Clr fused to the T25 or T18 domains of *Bordetella pertussis* adenylate cyclase 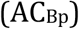 were produced in *E. coli* BTH101 by cloning the *cyaC* and *clr* genes, respectively, up- or down-stream of the gene fragments for T18 and T25. For *cyaC* variants CyaC# and CyaC## were used (Klein et al. 2023) that produce enzymatically inactive forms of CyaC. Use of the inactive variants of CyaC allows application of the BACTH system which relies on the activity of 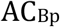 as the reporter and avoids interference from background cyclase activity of CyaC (Klein et al. 2023). CyaC# and CyaC## represent variants CyaC(K410A T484A) and CyaC(R495A N491A), where the catalytic ATP site or the stabilization site for the transition state of the dimer in catalysis are inactivated (Wissig et al. 2019; Tesmer et al. 1997; Linder & Schultz, 2003). Derivatives CyaC# and CyaC## produced in the BACTH test strain (Fig. 2, and Klein et al. 2023) very low background AC (and consequently β-galactosidase) activity of below 9 % for CyaC# (Klein et al. 2023) and CyaC## (Fig. 2) compared to the positive Zip-Zip control. The low background activity of the variants allowed their use for testing the restoration of *B. pertussis* 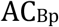 adenylate cyclase activity in the reporter strain. Strains producing CyaC#, or CyaC##, together with Smc01817 restored high (86 % of the positive control) or very high activity (152 % of the positive control). The data therefore indicates the restoration of high AC activity from T18 and T25 domains as a result of the CyaC# - Smc01817or CyaC## – Smc01817 provoked interactions.

**Figure 1:**
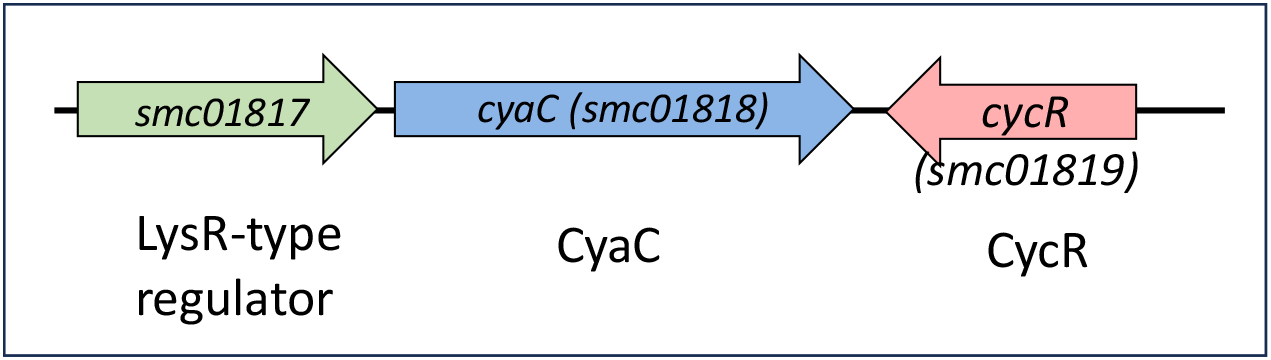
Gene arrangement around *cyaC* (*smc01818*) with *cycR* (*smc01819*) and *smc01817*. Orientation and relative lengths of the genes, as well as gene and protein names are indicated.

**Figure 2:**
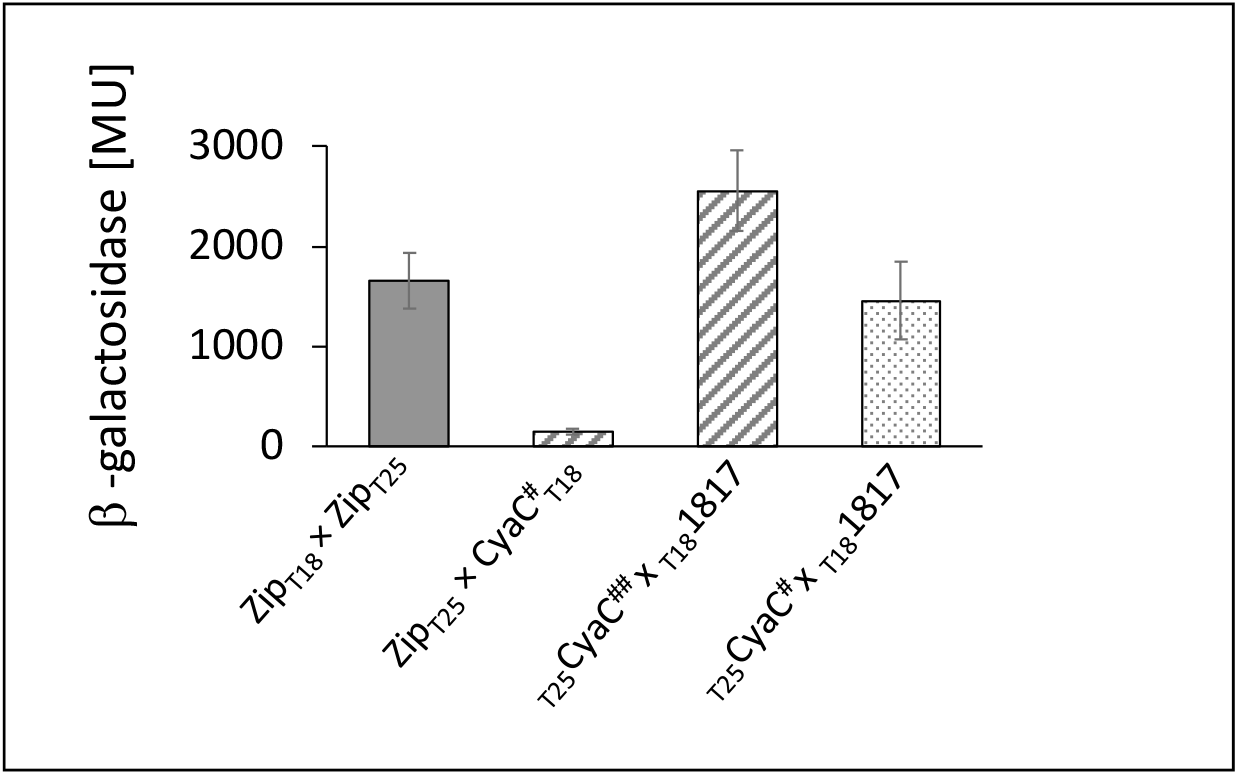
Interaction of CyaC and Smc01817 tested by the BACTH system for the pairs CyaC## / Smc01817 and CyaC# /Smc01817 in *E. coli* BTH101. Proteins CyaC# (Cyac(K410A T484A), CyaC## (Cyac(R495A N491A)and Smc01817, respectively, were fused C- or N-terminally to the T25 or T18 fragments of ACBP, as indicated. The proteins were produced pairwise from the corresponding genes from plasmids (Table 1). The Leu zipper (Zip) pair fused to T18 and T25 fragments, respectively, was used as the positive control, the pair Zip-T25 with CyaC#-T18 as the negative control. 100 % activity corresponds to 1,640 Miller-Units. Bacteria were grown anaerobically in LB medium supplemented with 20 mM dimethyl sulfoxide (Klein et al. 2013; Wissig et al. 2019). β-Galactosidase activities are given in Miller-Units MU (Miller 1992) as the mean (with standard deviation) from at least two biological and four technical replicates each.

### Interaction of Clr with Smc01817

In the CyaC×CycR×Clr complex the specific regulator CycR shows interaction with both the AC and the general regulator Clr (Klein et al. 2023). Therefore, here the interaction of Smc01817 with Clr was tested by the BACTH assay (Fig. 3). Smc01817 and Clr were used in the test in all suitable (eight) combinations of T18 and T25 fusions. Seven of the eight pairs showed high activities (700 to 1100 MU) which are well beyond 50% of the positive control (1370 MU). Only one of the pairs is with an activity of 460 MU which is below 50% of the positive control. The consistently high values strongly suggest therefore interaction of Smc01817 with Clr. Interaction of CyaC with Clr has been demonstrated earlier by different methods (Klein et al. 2023).

**Figure 3:**
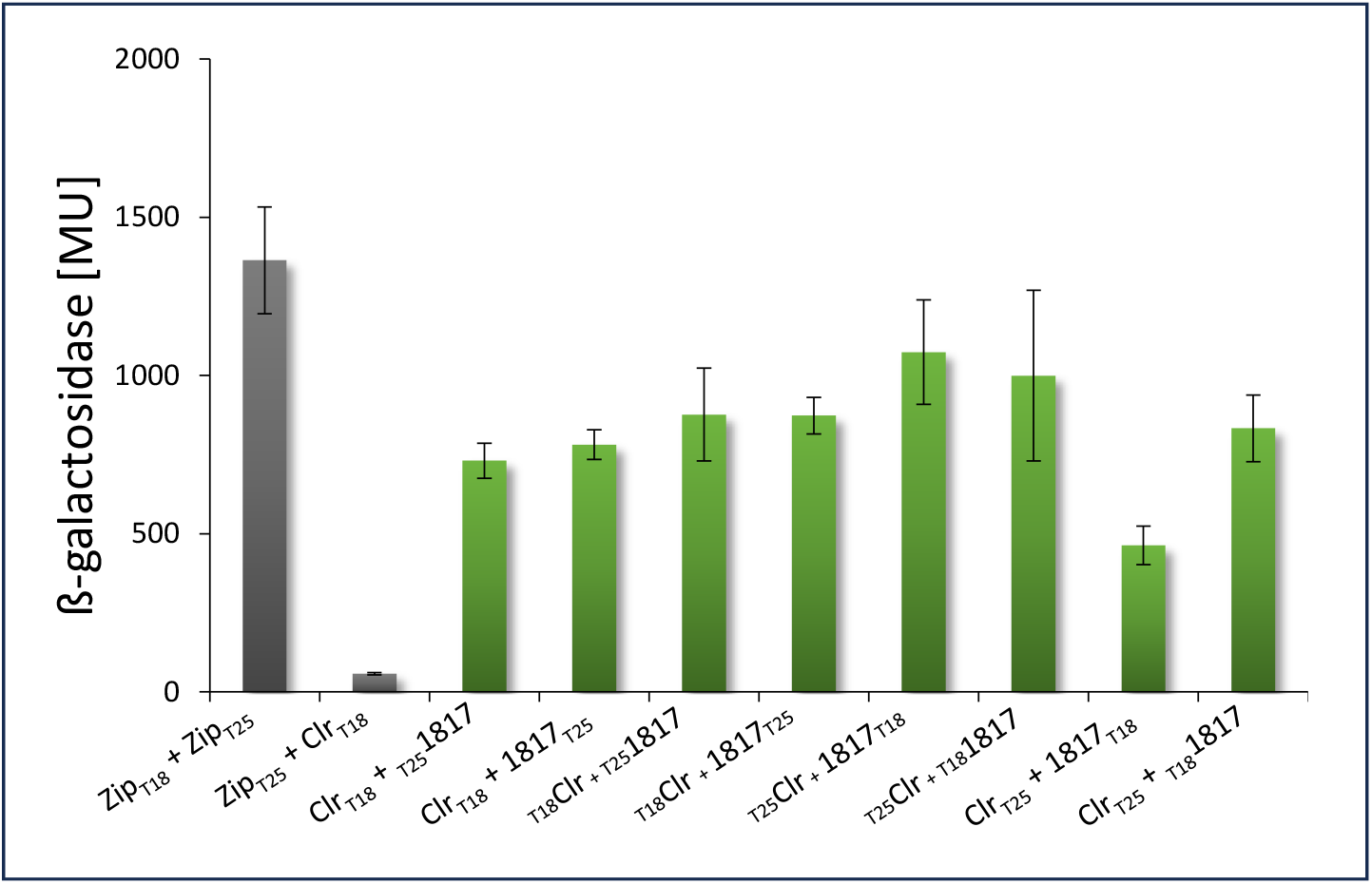
Interaction of Smc01817 with Clr tested by the Bacterial Two-Hybrid System (BACTH). Proteins Smc01817 (‘1817’) and Clr were fused C- or N-terminally to the T25 or T18 fragments of ACBP, respectively. The proteins were produced pairwise from the corresponding genes from plasmids (Table 1). The Leu zipper (Zip) pair fused to T18 and T25 fragments, respectively, was used as the positive control, the pair Zip-T25 with Clr-T18 as the negative control. 100 % activity corresponds to 1,370 Miller-Units. Bacteria were grown anaerobically in LB medium supplemented with 20 mM dimethyl sulfoxide (Klein et al. 2013; Wissig et al. 2019). β-Galactosidase activities are given in Miller-Units MU (Miller 1992) as the mean (with standard deviation) from at least two biological and four technical replicates each.

The interaction of the individual proteins CyaC×Smc01817 and Clr×Smc01817 appears to be specific since Smc01817 showed no interaction (< 5% of the positive control) with unrelated transcriptional regulators such as NreC (Fig. 4), similar to CyaC and Clr as demonstrated earlier (Klein et al. 2023).

**Figure 4:**
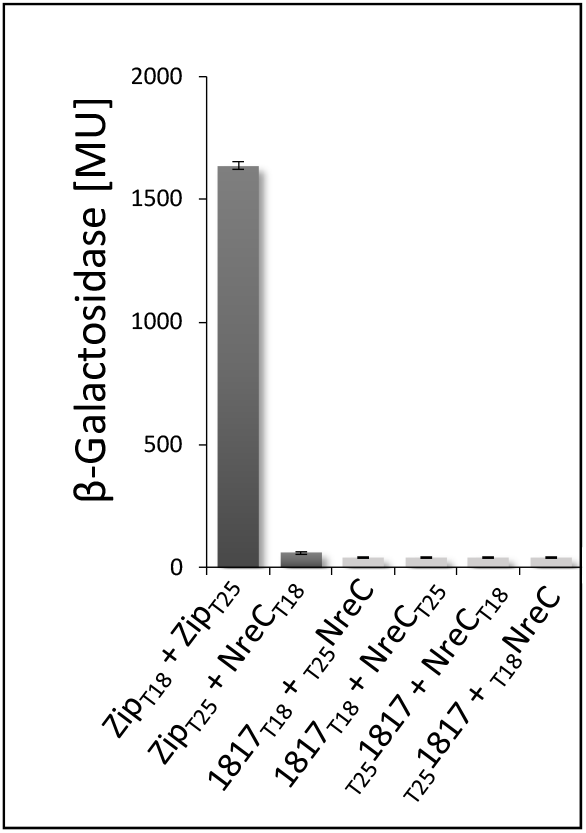
Interaction of Smc01817 with NreC as a cytosolic control protein tested by the Bacterial Two-Hybrid System (BACTH). Proteins Smc01817 (‘1817’) and NreC were fused C- or N-terminally to the T25 or T18 fragments of ACBP, respectively, as indicated. The proteins were produced pairwise from the corresponding genes from plasmids (Table 1). The Leu zipper (Zip) pair fused to T18 and T25 fragments, respectively, served as the positive control, the pair ZipT25 with NreCT18 as the negative control. Bacteria were grown anaerobically in LB medium supplemented with 20 mM dimethyl sulfoxide (Klein et al. 2013; Wissig et al. 2019). β-Galactosidase activities are given in Miller-Units MU (Miller 1992) as the mean (with standard deviation) from at least two biological and four technical replicates each.

## Discussion

### Specific signalling complexes with ‘caged’ cAMP vs freely diffusing cAMP as secondary messenger

The BACTH data demonstrate strong interaction of the LysR-type transcriptional regulator Smc01817 with adenylate cyclase CyaC and the cAMP binding transcriptional regulator Clr. Earlier it has already been shown by *in vivo* and *in vitro* experiments that CyaC interacts with Clr in an ATP (and thus cAMP) stimulated mode (Klein et al. 2023) (Fig. 5). Therefore, formation of a trimeric CyaC×Smc01817×cAMP-Clr complex is suggested, similar to the CyaC×CycR×cAMP-Clr complex that was demonstrated by *in vivo* interaction and *in vitro* data with purified proteins. Therefore, CyaC×Smc01817×cAMP-Clr is a supposed sensor-regulator complex with caged cAMP, for specific signalling from the sensory adenylate cyclase to target genes.

**Figure 5:**
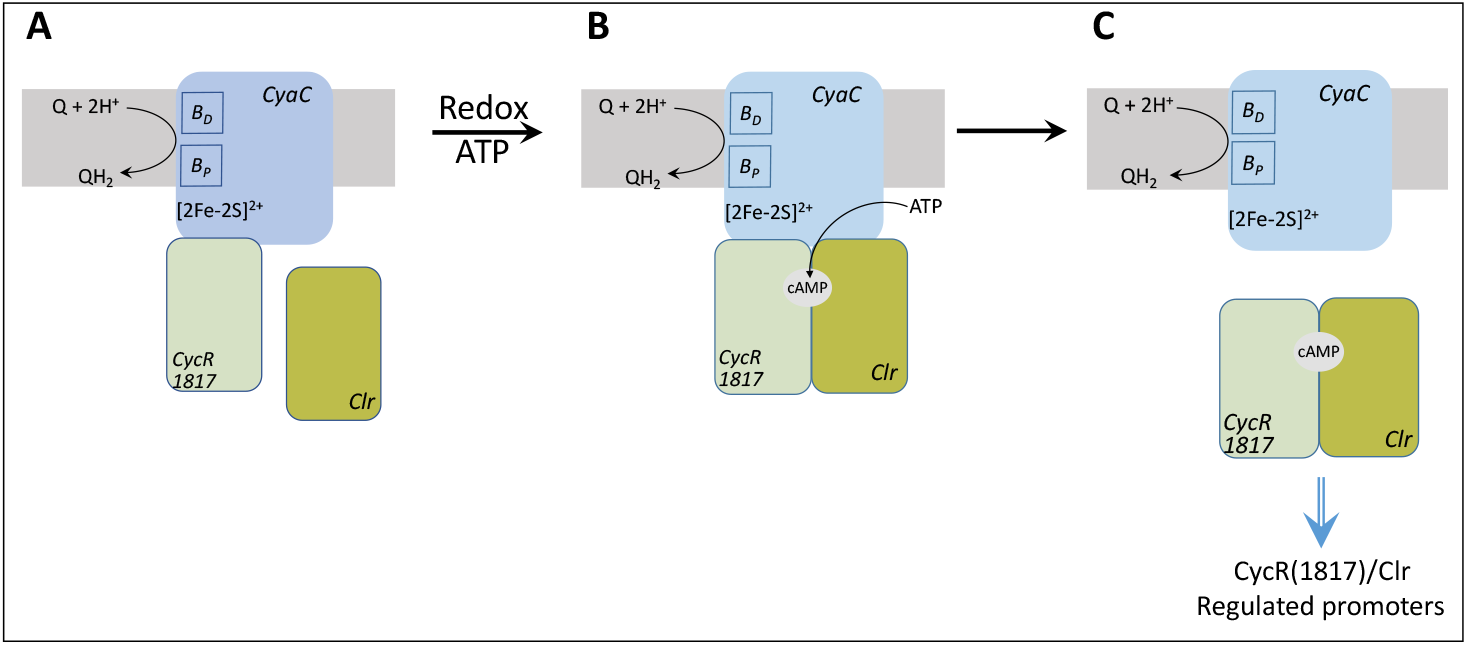
TheCyaC×CyaR×cAMP-Clr(and CyaC×Smc01817×cAMP-Clr) complexes and their function in signaling. (A) The AC CyaC forms with CycR the core of the complex at the membrane; interaction CyaC×CycR is independent of ATP and cAMP. (B) After redox activation, cyclase CyaC is active and forms from ATP cAMP. The cAMP is bound by Clr, and the trimeric CyaC×CycR×cAMP-Clr complex is formed which encloses the cAMP. (C) The CycR×cAMP-Clr complex is released to bind to CycR-Clr regulated promoters. A similar mechanism is suggested for CyaC×Smc01817×cAMP-Clr complex.

CycA×CycR×cAMP-Clr was supposed to control the expression of a subset of genes of the large Clr regulon of *S. meliloti* (Klein et al. 2023; Krol et al. 2016; Capela et al. 2001; Zou et al. 2017), that selects the subset of genes by the combined function of the specific regulator CycR and the general cAMP regulator Clr. The CycR×Clr were supposed to become specifically regulated after activation and cAMP production by CyaC, with the cAMP stored in the regulatory CyaC×cAMP-Clr complex (Klein et al. 2023). This model can be extended by the present work to suggest the LysR-type regulator Smc01817 forms the CyaC×Smc01817×cAMP-Clr complex for the regulation of another subset of genes of the Clr regulon that are also part of the CyaC responsive genes. Dual transcriptional control of genes by promoter regions that respond to an overriding and a specific regulator is common and implemented in various forms. This can be performed by regulators interacting only at the promoter region, such as catabolite control by CRP of *E. coli* together with the lactose inducer LacI at the *lacZYA* operon (Reznikoff 1992), or by the oxygen sensor FNR exerting co-dependent regulation of the *nirB* operon together with the nitrate regulator NarL (Tyson et al. 1993; Wu et al. 1998; Busby 2019; Beiner and Kiley 1999). CRP and LacI, FNR and NarL, respectively, cooperate at the promoters but are independent proteins, whereas cAMP-Clr and CycR (Klein et al. 2023), and presumably cAMP-Clr and Smc01817 are organized in a complex. In the NarL-DevR system for nitrate regulation in *Mycobacterium tuberculosis*, however, the nitrate regulator NarL becomes only active when it forms a heterodimer with the activated form of the DNA-binding protein DevR (Malhotra et al. 2015), resembling in this respect the supposed function of CycR×cAMP-Clr or Smc01817×cAMP-Clr, respectively.

Finding a second signalling complex consisting of an adenylate cyclase with transcriptional regulators, including an cAMP binding regulator, suggests that signalling complexes with ‘caged’ cAMP are commonly formed by adenylate cyclases in *S. meliloti*. This set-up avoids freely diffusing cAMP and allows specific signalling when small diffusing molecules are used by multiple signalling systems. Here, the cAMP is formed within the complex and serves as a trigger for activating the complex, rather than a diffusing secondary messenger in its classical form.

### Specific signalling in multiple c-di-GMP regulated systems

c-di-GMP represents another secondary messenger with broad distribution in bacteria. It controls numerous aspects of bacterial growth and behavior such as motility, biofilm formation, development and progression of the cell cycle, sporulation, aggregation and surface adhesion, and virulence of animal and plant pathogens, and more than 100 systems can respond to c-di-GMP within one bacterial species (D’Argenio 2004; Jenal et al. 2006; Römling et al. 2005: Hengge 2010b). Four major principles have been identified to control signalling specificity (Kunz & Graumann 2020). First, ‘global’ c-di-GMP signaling hypothesis A and B assume basically similar c-di-GMP levels in the cell. In hypothesis A the classifying principle is based on a difference in the binding affinity of c-di-GMP to the different effector (or receptor) proteins. In *Salmonella enterica* or Pseudomonas aeruginosa the binding affinities of the c-di-GMP sites show differences in binding affinities of up to 140-fold (Pultz et al. 2012). In hypothesis B signal specificity is based on the temporal separation of formation or activation of the receptor proteins, such as different time points for their expression during cell growth.

The ‘local’ c-di-GMP signaling hypothesis suggests two different modes (C and D). Hypothesis C assume direct interactions between the protein components DGCs (diguanylate cyclases)s, PDEs (phosphodiesterases) and the c-di-GMP receptors (effectors) which has been shown for systems in *B. subtilis* (Bedrunka and Graumann 2017, Kunz et al. 2020), *E. coli* (Lindenberg et al. 2013), *Pseudomonas* (Dahlstrom et al. 2015) and others. Thus, in *E. coli* the diguanylate cyclase has been shown to interact with phosphodiesterase YciR in the c-di-GMP signaling cascade for biofilm synthesis control. In the local c-di-GMP signalling of type C, proteins interact directly and set up a network or signaling pathways to control downstream signaling cascades for the control of cellular processes.

The second mode (type D) of ‘local’ c-di-GMP signaling assumes close local proximity of the proteins in c-di-GMP signalling in cellular compartments (subcellular clustering) without the direct need for protein complex formation. The first known system of this type was the chemotaxis Wsp system of c-di-GMP signaling in *P. aeruginosa*, which forms subcellular clusters (Güvener et al. 2007).

## Acknowledgements

We are grateful to Deutsche Forschungsgemeinschaft DFG and Johannes Gutenberg University of Mainz for supporting of the work

